# Identifying Women at Risk for Polycystic Ovary Syndrome Using a Mobile Health Application

**DOI:** 10.1101/659318

**Authors:** Erika Rodriguez, Daniel Thomas, Anna Druet, Marija Vlajic Wheeler, Kevin Lane, Shruthi Mahalingaiah

**Affiliations:** Department of Obstetrics and Gynecology, Boston University School of Medicine, 85 East Concord Street, Boston MA 02118, USA; Department of Epidemiology, Boston University School of Public Health, Talbot 3E, 715 Albany Street, Boston, MA 02118, USA; Clue®, BioWink GmbH, Adalbertstraße 7-8, 10999 Berlin, Germany; Department of Environmental Health, Boston University School of Public Health, Talbot T4W, 715 Albany Street, Boston, MA 02118; Department of Physiology & Biophysics, Boston University School of Medicine, 700 Albany St. W302, Boston, MA 02118, USA

**Keywords:** Polycystic Ovary Syndrome, mobile health app, Clue®, menstrual irregularities, telemedicine

## Abstract

**Background:** Polycystic ovary syndrome (PCOS) is an endocrine disrupting disorder affecting at least 10 percent of reproductive-aged women. Women with PCOS are at increased risk for diabetes and cardiovascular disease. In North America and Europe, the diagnosis of PCOS may be delayed several years and may require multiple doctors resulting in lost time for risk-reducing interventions. Menstrual tracking applications are one potential tool to alert women of their risk for PCOS while also prompting them to seek evaluation from a medical professional.

**Objective:** The objective of this study was to develop the *Irregular Cycles Feature* (ICF), an adaptive questionnaire, on the mobile phone application (app) Clue® to generate a probability of a virtual test subject’s risk for PCOS. The secondary objective was to assess the accuracy of the ICF by comparing the probability of risk generated by the app to a probability generated by a physician.

**Methods:** First, a literature review was conducted to generate a list of signs and symptoms of PCOS, termed variables. These include, but are not limited to, hirsutism, acne, and alopecia. Probabilities were assigned to each variable and built into a Bayesian network. The network served as the backbone of the ICF, which identified potential subjects through self-reported menstrual cycles and answers to medical history questions. Upon completion of the questionnaire, a Result Screen summarizing the virtual test subject’s probability of having PCOS is displayed. For each eligible virtual test subject, a Doctor’s Report containing information regarding tracked menstrual cycles and self-reported medical history is generated. Both of these documents share information about PCOS and detailed explanations for facilitating a diagnosis by a medical provider. Virtual test subjects were assigned probabilities by a) the ICF and b) a board-certified reproductive endocrinology/infertility physician-scientist, which served as the gold standard. The ICF was set to recommend individuals with a score greater than or equal to 25% to follow-up with their physician. Differences between the network and physician probability scores were assessed using a t-test and a Pearson correlation coefficient. An additional iteration was performed to improve the ICF’s prediction capability.

**Results:** The first iteration of the ICF produced only one false positive compared to the physician screening score and had an absolute mean difference of 15.5% (SD= 15.1%) amongst virtual test subjects. Upon modification of the ICF, the second iteration had two false positives as compared to the physician screening score and had an absolute mean difference of 18.8% (SD = 13.6%). The majority of virtual test subjects had an ICF score that over predicted PCOS when compared to the physician. However, there was strong positive significant correlation between the ICF and the physician score (Pearson correlation coefficient= 0.69; p < 0.01). The second iteration performed worse with a Pearson correlation coefficient of 0.54; p > 0.01).

**Conclusion:** The first iteration ICF, as compared to the second, was better able to predict the probability of PCOS and can potentially be used as a screening tool to prompt a high-risk subject to seek evaluation by a medical professional.

## Introduction

According to the Rotterdam criteria, polycystic ovary syndrome (PCOS) is clinically diagnosed by the presence of at least 2 of the following: androgen excess, menstrual irregularity, or presence of polycystic ovary morphology upon ultrasound examination [1]. The Androgen Excess and PCOS Society has also proposed guidelines for diagnosis which include: hyperandrogenism, ovarian dysfunction, and exclusion of other androgen excess or related disorders [2]. Women with menstrual irregularities, particularly those with PCOS, have an increased risk of developing comorbidities, such as metabolic syndrome, heart disease or diabetes [3]. In 2005, the economic burden of evaluating and providing care to women of reproductive age with PCOS was $4.36 billion United States dollars (USD), which is the equivalent of $5.65 billion USD in 2018. This assessment included the costs of infertility treatments, living and treating metabolic disorders, and addressing hirsutism. The calculated expenses of PCOS are likely an underestimate of the actual cost of providing care because many women will live beyond reproductive age with expensive metabolic disorders [4].

In the current era of ubiquitous smartphones, individuals are turning to mobile health applications (apps) for immediate health tracking and care [5, 6]. Women in particular are reporting higher rates of downloading health apps [7]. By having menstrual cycle data collected in real time, women are able to share accurate details with their medical providers. Thus, these apps can have significant implications for women with menstrual irregularities.

For women at high risk of developing PCOS, having a mobile health app that identifies and tracks menstrual cycles is crucial. The objective was to develop the Irregular Cycles Feature (ICF) to flag menstrual cycles that are clinically out-of-range while capturing relevant symptoms to generate health summaries for women with menstrual irregularities.

## Materials and Methods

Clue® is an app, created by the company Bio Wink Gmbh, that allows users to track menstrual cycles and related health information. Data collected includes, but is not limited to, menstrual cycle length, duration of flow, menstruation related pain symptoms, and method of birth control. In order to create a user profile, the app also prompts individuals to input age, height, and weight.

An adaptive questionnaire, or a question set that is modified based on answers to previous questions, was developed. To create the framework for the adaptive questionnaire, the board-certified reproductive endocrinology/infertility physician-scientist (physician) from Boston University and researchers at Clue® compiled signs and symptoms (variables) associated with PCOS. A literature review was then conducted in order to determine the prevalence of these variables specifically in women with PCOS. Each variable and associated probability was then incorporated into a Bayesian network (Figure 1).

**Figure 1.**
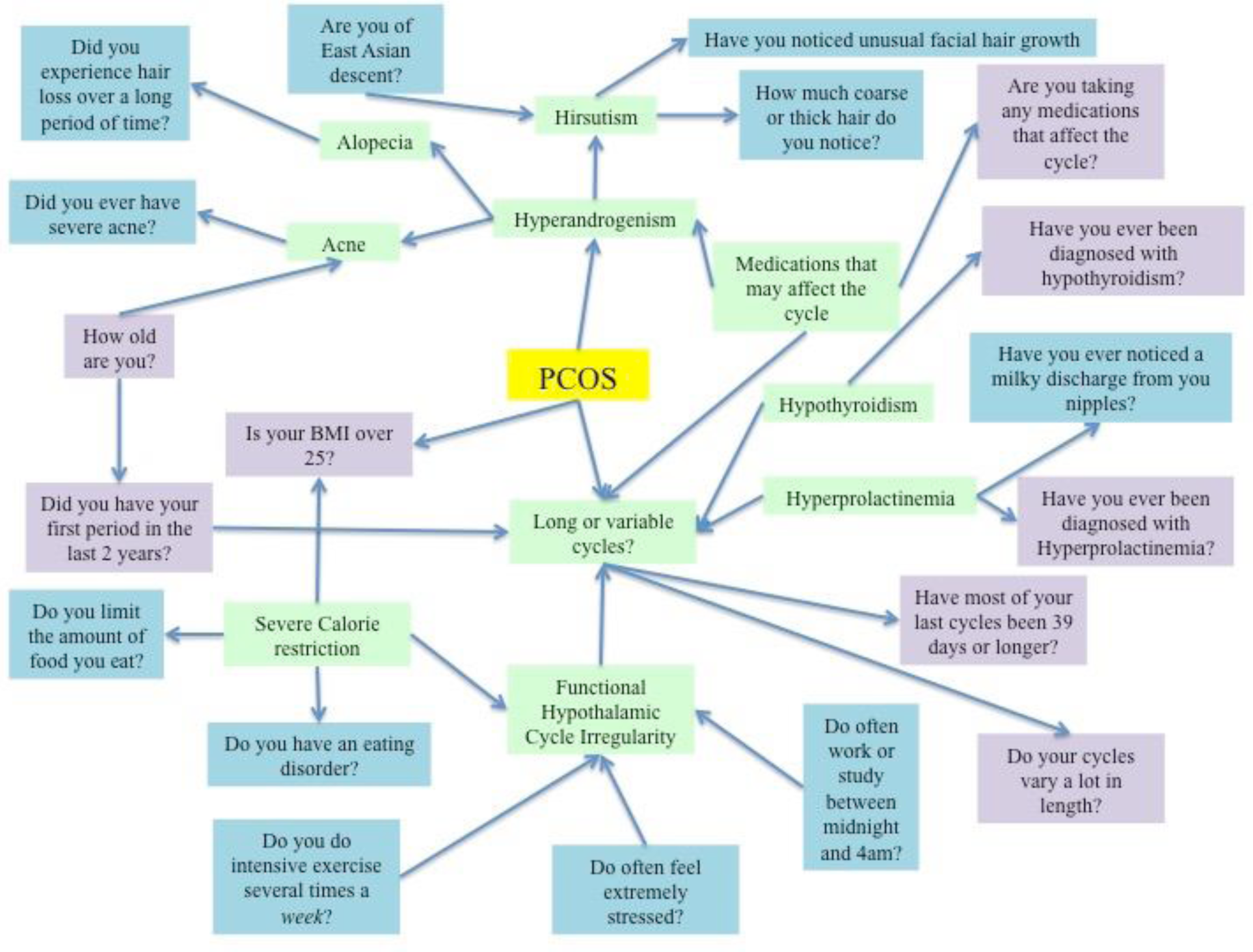
Green boxes indicate major diagnostic concerns for PCOS. Purple boxes are representative of the questions that are in the Screener. Blue boxes are additional questions asked by the ICF.

### Phase One: Development of the Question Sets and Virtual Test Subjects

Once the network was established, questions were written to assess signs and symptoms deemed relevant by the literature review. Questions were separated into a Screener and the adaptive questionnaire, the basis of the ICF. The Screener consists of eight questions used to collect information regarding height, weight, age, birth control use, medical conditions, life stage, age at menarche, and cycle length.

After the Screener, the subject is presented with the adaptive questionnaire, which features questions aiming to complete the picture of risk factors for PCOS. These questions are based on the Ovulation and Menstruation Health Study conducted at Boston University, which measures androgen excess, stress, and eating patterns. Questions in the adaptive survey include: measures of body hair, medications consumed that can affect the menstrual cycle, hair loss, acne, irregular sleep, stress, strenuous physical work, and eating habits. While this is not an exhaustive medical history, these variables were deemed useful for preliminary identification of PCOS based on the current literature. A complete list of additional questions assessed by the feature can be seen in Table 1. Because some symptoms vary by racial and ethnic groups, users are also given the option to report their background.

**Table 1.** Categories to assess menstrual irregularities. Three different categories for assessing risk of PCOS were established based on the Ovulation and Menstruation Health Study. Each of the questions in the ICF questionnaire (excluding the Screener) is included in the table.

|  | **Assessing Androgen Excess** | **Assessing Lifestyle** | **Assessing Health** |
| --- | --- | --- | --- |
| **Question 1** | Which categories describe you? Select all that apply | Do you often feel extremely stressed, or do you work in a very high-stress job? | Are you taking any medications that may affect your hormones or menstrual cycle? Or have you stopped using this medication within the past 3 months? (This could be hormonal birth control, hormone therapy, testosterone, etc.) |
| **Question 2** | Are you concerned about the amount of coarse or thick hair that grows on your face and/or body (including on your upper lip, chin, face, chest, back, abdomen, arms, and thighs)? | Do you often work or study at least some time between midnight and 4:00AM? | Do you often limit the amount of food you eat to influence your shape or weight, or to relieve stress? |
| **Question 3** | How much coarse or thick hair do you notice on these areas? | Do you exercise 5 days a week or more, for at least an hour each time? | Have you noticed a milky discharge coming from your nipples (outside of recent pregnancy or breastfeeding)? |
| **Question 4** | Do you notice that the hair on your head is thinning faster than other people your age? |  |  |
| **Question 5** | Do you have acne that you consider to be severe |  |  |

Fourteen virtual test subjects were put through the Screener in order to assess the functionality of the Bayesian network. Each of the virtual test subjects had unique answers to the ICF Screener, and then questionnaire, in order to test the ability of the network to produce an accurate PCOS probability. Each virtual test subject had certain questions left blank, in order to test the accuracy of the results generated in case a user chose not to answer certain questions. Of the 14 virtual test subjects that were screened, 9 were deemed eligible to continue onto the ICF in order to ascertain if PCOS is a possible cause of their menstrual irregularities.

### Phase Two: Question Flow

Because the ICF is adaptive, not every question is asked to every subject. For instance, Figure 2 shows that if a user reports that they are not concerned about excess hair growth, the tool will not ask about how much hair is present. In this manner, the model streamlines the data collection and reduces the time it takes an individual to complete the module. Compiling this information, the network can then calculate whether menstrual cycle irregularities are an indicator for PCOS or if the menstrual irregularities are possibly due to a different condition.

**Figure 2.**
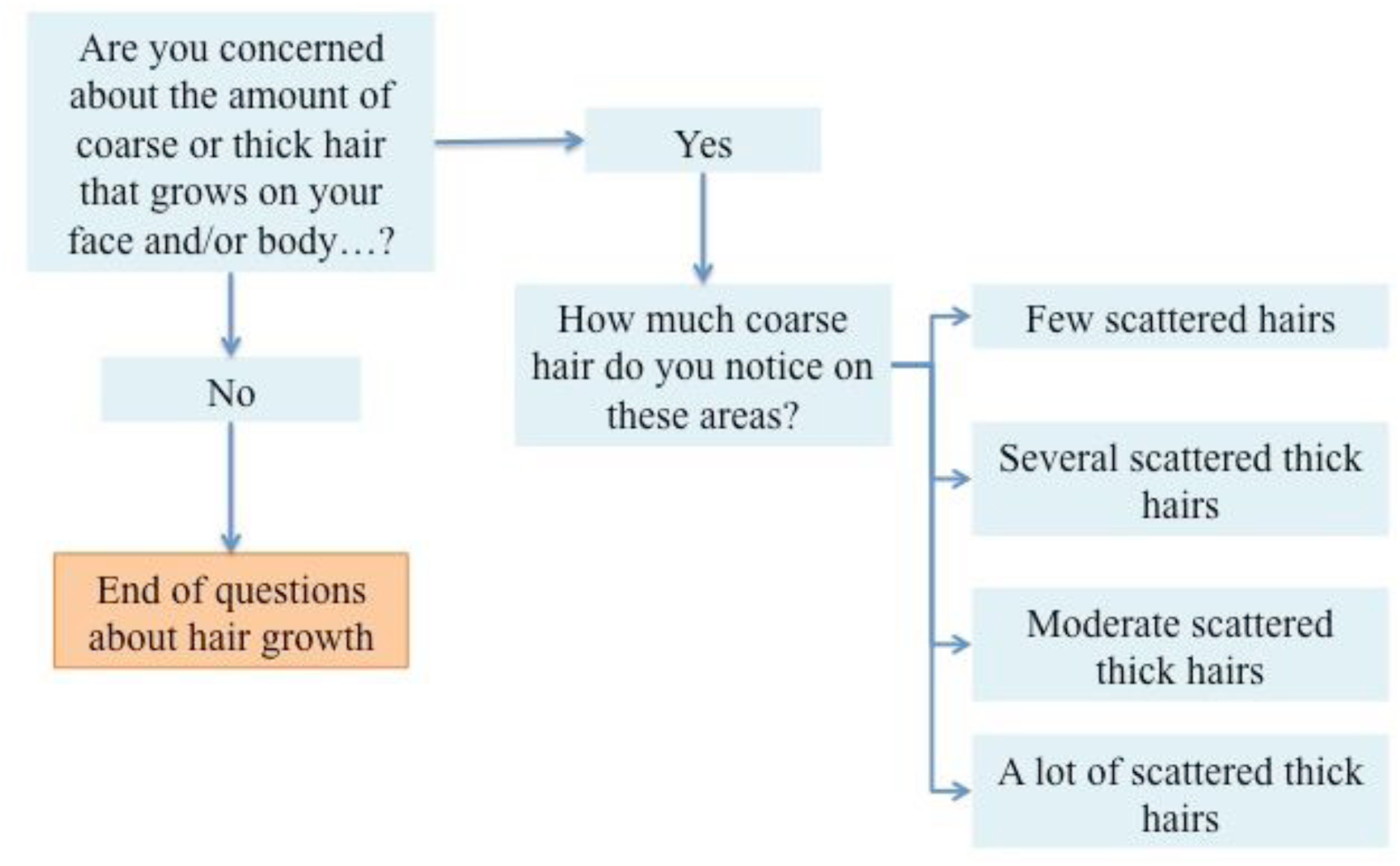
This figure highlights the adaptive nature of the ICF by modeling with a questions assess hair growth. The orange box indicates the end of the question set.

### Phase Three: Assignment of Thresholds for Risk

The physician on the team assigned four categories of risk: low, indeterminate, moderate, or high to each virtual test subject. In order to compare this to the quantitative values generated by the ICF, the team determined a set of numerical ranges for the categories based on proposed definitions for converting between qualitative and quantitative classifications by Hillson in 2005 [8].

Virtual test subjects determined to have low risk of PCOS were assigned a value within the range of 0% to 9%. The numerical range for indeterminate risk was assigned as 10% to 29%. These values were selected based on the ‘unlikely’ and ‘possible’ qualitative terms in Hillson [8]. Virtual test subjects with moderate individual risk were assigned percentages within the range of 30% to 59%. These values encompassed the categories of ‘probable’ and ‘a good chance’ as defined by Hillson [8]. Virtual test subjects with high individual risk were assigned a value within the range of 60-100%. These values were selected based on the Hillson’s interpretation of ‘highly probable’ to ‘definite’ terms [8]. These probability scales are summarized in Figure 3.

**Figure 3.**
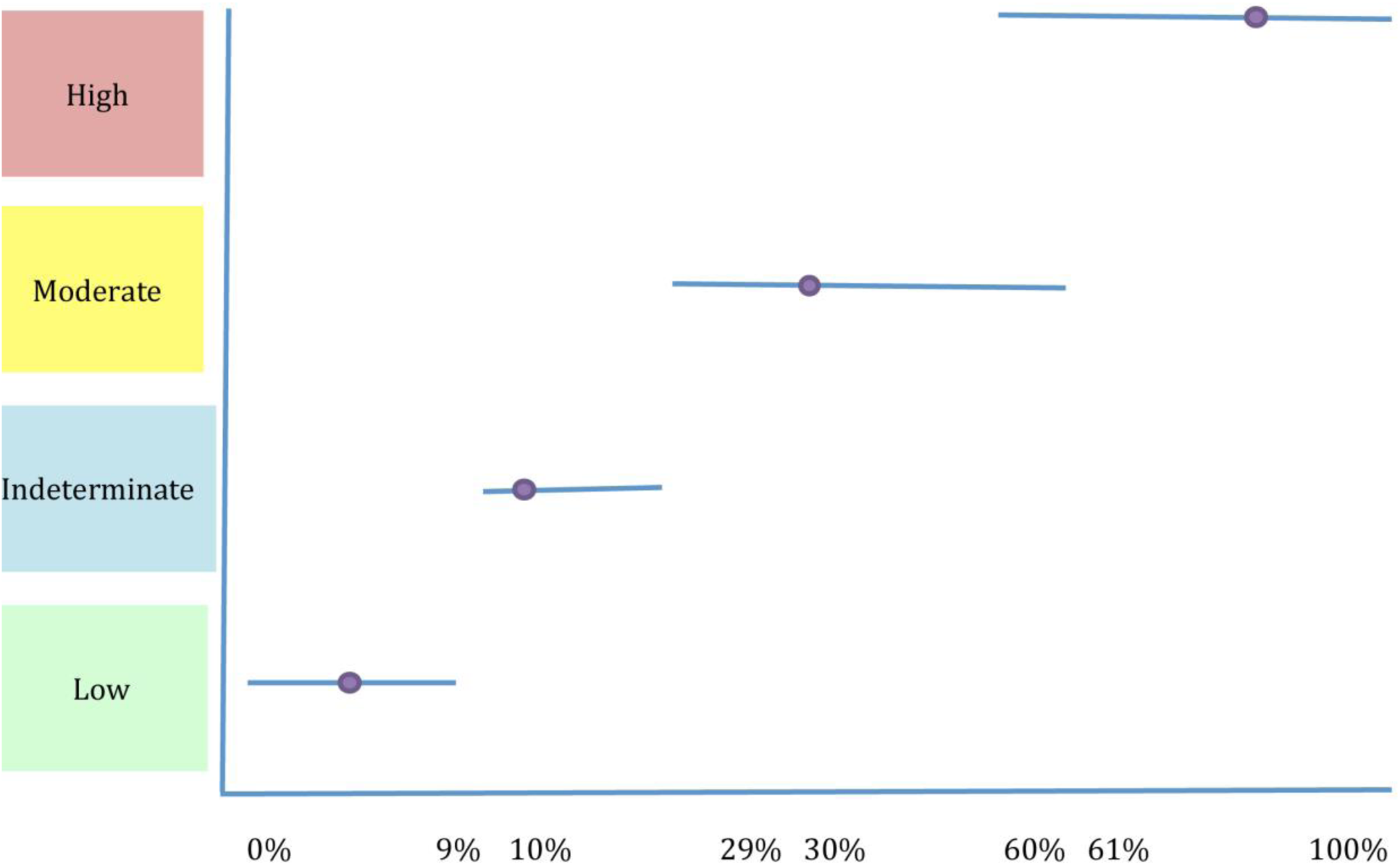
Sliding Scale Percent Probability Ranges: Adapted from Hillson, 2005. The blue bar indicates the range of percentages that fall into the high, moderate, indeterminate, and low categories. The purple circle illustrates the percentage typically used in assignment of percent probability.

During the evaluation of the virtual test subjects, the physician determined that two virtual test subjects were confounded by additional variables and could not be accurately assessed due to lack of data. These two cases were assigned a probability of 10% as they fell into this indeterminate category. 10% was arbitrarily chosen for this study because it is reflective of a low but possible risk of developing PCOS.

Results Screens were created to provide feedback at the end of the assessment based on each of these thresholds. Distinct and direct recommendations are given, based on the calculated probability of PCOS yielded by the ICF. There are three possible screens including a Positive Result Screen, a Neutral Result Screen, and an Inconclusive Result Screen. All screens include a brief description of how the result was reached, and a disclaimer that the result is not a diagnosis.

Users calculated as having a probability of PCOS that is greater than or equal to 25% by the ICF are prompted with a Positive Result Screen (Supplemental Figure 1). This screen displays: a description of PCOS, related health risks, and a call to action encouraging the user to seek medical attention. Additionally, it describes the steps a doctor may take in order to diagnose the individual with PCOS or another related disorder affecting menstrual cycle regularity.

The Neutral Result Screen is presented to users with an ICF probability of less than 25%. It states that a prediction cannot be made regarding what is causing the irregularities. It includes potential other causes such as lifestyle factors, hypothyroidism, and Cushing’s syndrome. The user is also prompted to seek advice from a medical professional. The text describes that the doctor will likely perform a detailed history regarding symptoms, a simple physical exam, and blood tests if necessary.

The Inconclusive Screen is presented to users who have reported confounding variables or too much missing data. These variables include, but are not limited to: the use or recent discontinuation of hormonal birth control, age outside applicable range, recent pregnancy, and breastfeeding. The potential use of medications, particularly those with hormones, in individuals creates too many confounding variables and is currently beyond the capabilities of the network’s calculations. Thus, when an individual reports a confounding variable that also causes hormonal dysregulation, the screen prompts them with a suggestion to visit a medical professional who can do additional testing to determine the reason behind their menstrual irregularity. Six of the 14 virtual test subjects were ultimately prompted with an Inconclusive Screen due to their answers.

The Doctor’s Report is a shareable document that is generated at the end of each assessment for presentation to a medical professional (Supplemental Figure 2). It includes details regarding the user’s menstrual cycle characteristics and history, as well as the signs and symptoms reported via the ICF questionnaire portion. The medical board at Clue® and the physician from Boston University were consulted in its design in order to ensure that it appropriately highlighted a user’s health data. The Doctor’s Report is generated whether the individual is expected to have PCOS or not. This ensures that the individual is able to provide important medical information to their provider regardless of the output so long as their cycles are irregular.

### Phase 4: Validation of the Tool

In order to validate the usability and accuracy of the network, the physician was asked to determine whether or not the probabilities predicted by the ICF were similar to the assessments she would make in a clinical setting. A summary of the probability assignments by both the ICF and the physician can be seen in Table 2.

**Table 2.** Probability assignments. A summary of the physician assessments (categorical probabilities), physician’s assigned probabilities, and the ICF calculated probability for all test subjects that went through the ICF questionnaire during the first iteration.

| **Subject Number** | **Physician Assessment** | **Physician Assigned Probability** | **ICF Assigned Probability** |
| --- | --- | --- | --- |
| **Subject 1** | High | 80% | 91% |
| **Subject 2** | High | 80% | 93% |
| **Subject 8** | PCOS: 30%<br>Hypothalamic pituitary: 70% | 30% | 66% |
| **Subject 12** | Currently taking hormones:<br>This is a special case-<br>indication of irregular cycles<br>means the feature should<br>direct them to talk to their<br>doctor | 10% | 10% |
| **Subject 14** | Low PCOS | 5% | 34% |
| **Subject 17** | Moderate to high PCOS | 70% | 37% |
| **Subject 5** | Indeterminable | 10% | 9% |
| **Subject 7** | High | 80% | 14% |
| **Subject 15** | None | 0% | 1% |

### Statistical Analysis

The utility of the ICF was assessed by (1) comparing the sensitivity (false negatives) and specificity (false positives) to the physician assessment, (2) mean difference amongst virtual test subject scores, and (3) Pearson correlation coefficient with the physician score. The correlation between the ICF and physician, a p-value from the correlation, and a Pearson correlation coefficient were calculated using the 9 eligible virtual test subjects. A second analysis was then conducted excluding 1 virtual test subject because it significantly skewed the data, which resulted in a final set of 8 virtual test subjects that were used to assess the sensitivity and specificity, mean difference, and Pearson correlation coefficient.

An additional iteration of the ICF’s Bayesian network was also completed to determine the effect of hirsutism on a PCOS subject. This was done using the final set of 8 virtual test subjects. The modification was deemed necessary because current literature suggests lower probabilities than what was found in the original review. A summary of the statistical tests can be seen in Tables 3-6.

**Table 3.**
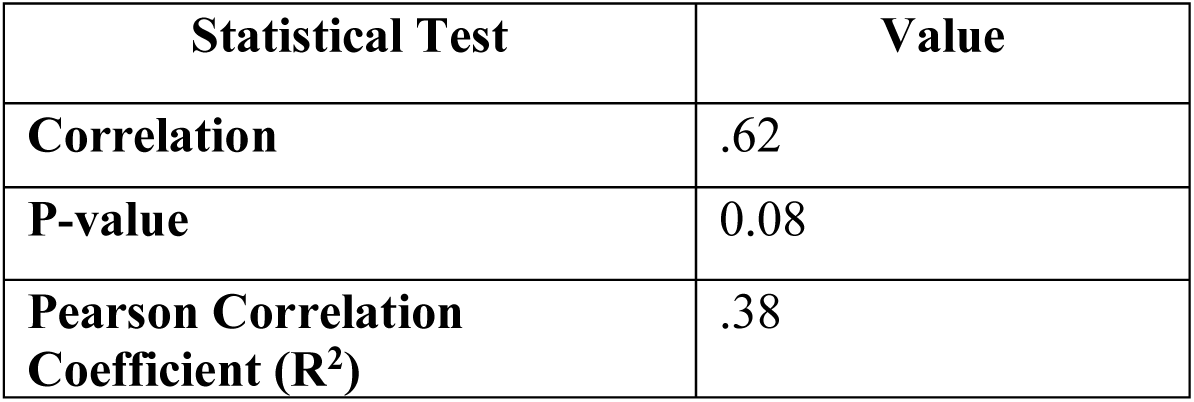
Summary of statistical calculations for 9 virtual test subjects. This table demonstrates the statistics for all virtual test subjects that were created by the team including subject 7 who was ultimately excluded because it was suspected to be an outlier based on the difference in statistical calculations.

| Statistical Test | Value |
| --- | --- |
| **Correlation** | .62 |
| **P-value** | 0.08 |
| **Pearson Correlation Coefficient ($R^2$)** | .38 |

**Table 4.**
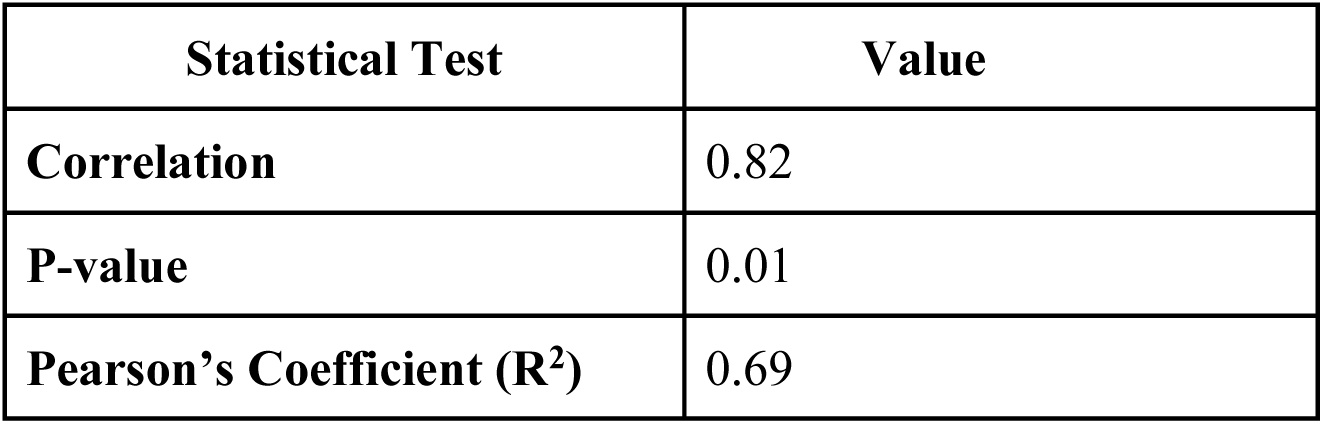
Summary of statistical calculations for 8 virtual test subjects. This table demonstrates the statistics for eight test subjects that were created by the team. It excludes subject 7 who was suspected to be an outlier based on the mean difference.

| Statistical Test | Value |
| --- | --- |
| **Correlation** | 0.82 |
| **P-value** | 0.01 |
| **Pearson's Coefficient ($R^2$)** | 0.69 |

**Table 5.** Summary of statistical calculations for 8 virtual test subjects on 2nd iteration of ICF. This table demonstrates the statistics calculated with lowered probabilities of hirsutism in PCOS for all test cases excluding subject 7 (the suspected outlier).

| Statistical Test | Value |
| --- | --- |
| **Correlation** | 0.73 |
| **P-value** | 0.03 |
| **Pearson's Coefficient ($R^2$)** | 0.54 |

**Table 6.** Sensitivity, specificity, and mean difference of ICF and Physician scores by iteration.

| Iteration | Patient | Network | Physician | Specificity | Sensitivity | Difference |
| --- | --- | --- | --- | --- | --- | --- |
| 1 | 1 | 91% | 80% | 0 | 0 | 11 |
|  | 2 | 93% | 80% | 0 | 0 | 13 |
|  | 8 | 66% | 30% | 0 | 0 | 36 |
|  | 12 | 10% | 10% | 0 | 0 | 0 |
|  | 14 | 34% | 5% | 1 | 0 | 29 |
|  | 17 | 37% | 70% | 0 | 0 | -33 |
|  | 5 | 9% | 10% | 0 | 0 | -1 |
|  | 15 | 1% | 0% | 0 | 0 | 1 |
| Mean Absolute Difference |  |  |  |  |  | 7 |
| 2 | 1 | 88% | 80% | 0 | 0 | 8 |
|  | 2 | 92% | 80% | 0 | 0 | 12 |
|  | 8 | 56% | 30% | 0 | 0 | 26 |
|  | 12 | 10% | 10% | 0 | 0 | 0 |
|  | 14 | 26% | 5% | 1 | 0 | 19 |
|  | 17 | 29% | 70% | 0 | 0 | -31 |
|  | 5 | 22% | 10% | 0 | 0 | 12 |
|  | 15 | 42% | 0% | 1 | 0 | 42 |
| Mean Absolute Difference |  |  |  |  |  | 11 |

## Results

The majority of virtual test subjects had an ICF score that over predicted PCOS as compared to the physician score (Table 6). The first iteration of the ICF produced only one false positive as compared to the physician screening score and had an absolute mean difference of 15.5% (SD = 15.1%) amongst virtual test subjects. The second iteration of the ICF had two false positives as compared to the physician screening score and had an absolute mean difference of 18.8% (SD= 13.6%). The first iteration ICF score over predicted the probability of PCOS as compared to the physician with a mean absolute difference of 7. A correlation value was calculated to be 0.82 for the 8 virtual test subjects. The same 8 virtual test subjects were also used to generate a linear regression. The Pearson correlation coefficient for this regression was calculated to be 0.69 (Figure 4). The p-value was then determined to be 0.01 (Table 4). Because the p-value is less than 0.05, the team determined the predictive capabilities of the ICF were statistically significantly different to the assessments made by the physician although the risk difference was not considered to be clinically significant. A few sample virtual test subjects are shown in Table 7 in order to demonstrate the questions, answers, and predictions that were generated by the ICF.

**Table 7.** Sample questions, answers, and probabilities using the ICF. This displays select virtual test subject answers to the ICF questionnaire. The questions in blue are Screener questions.

| Question | Subject 1 | Subject 8 | Subject 12 | Subject 15 |
| --- | --- | --- | --- | --- |
| Concerned about hair growth? | Yes | Yes | Yes | No |
| How much thick hair? | Lots of Hair | Several | Several |  |
| Majority of cycles $\geq 38$ days? | Yes | Yes | Yes | No |
| Taking meds that could affect the cycle? | No | No | Yes |  |
| BMI over 25 | No | No | No |  |
| Cycle variation out of range? | Yes |  |  | No |
| Eating disorder? | No | Yes |  |  |
| Hair loss? | No | No |  |  |
| Acne? | No | No |  |  |
| Menarche during last 2 years? | No |  |  |  |
| Age Range? |  | 19-40 |  |  |
| Of East Asian heritage? |  |  |  |  |
| Diagnosed with Hypothyroidism? |  |  |  |  |
| Irregular sleep? |  |  |  |  |
| Stress? |  |  |  |  |
| Strenuous physical work? |  |  |  |  |
| Limiting food? |  |  |  |  |
| Diagnosed with Hyperprolactinemia? |  |  |  |  |
| **PCOS Probability** | **91%** | **66%** | **10%** | **1%** |

**Figure 4.**
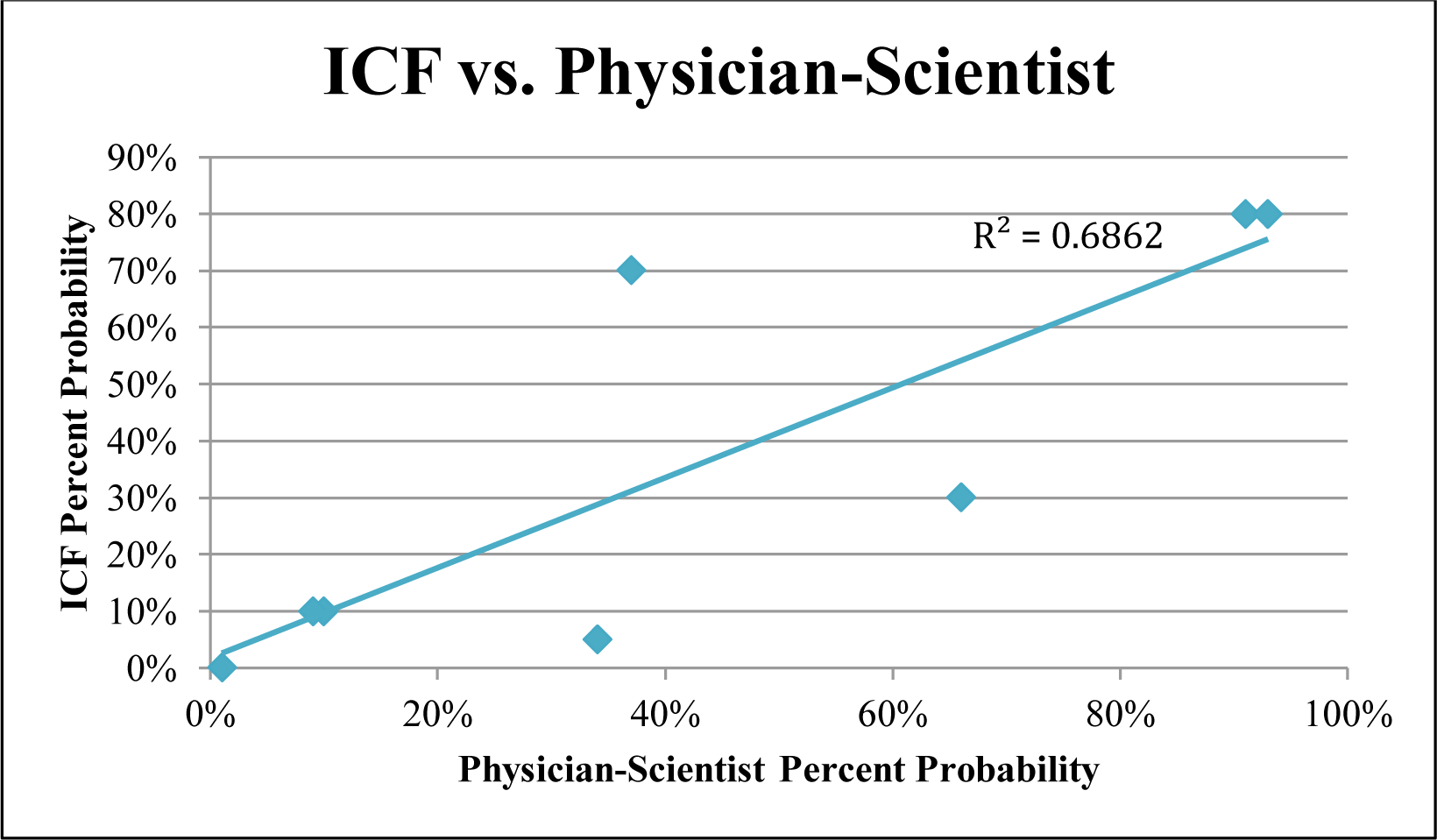
Linear regression with Pearson’s coefficient for 8 test cases. This graph demonstrates the linear regression for 8 test cases, which excludes subject 7 (the suspected outlier).

The results generated by the ICF as compared to those assigned by the physician can be seen in Table 2. For virtual test subject 1, the physician assigned an 80% probability, as compared to the 91% calculated by the ICF. In both cases, the user would be prompted with a Positive Result Screen and the Doctor’s Report reporting PCOS as a possible cause for the irregular menstrual cycles.

In virtual test subject 8, the physician assigned a 30% probability of PCOS, and further suggested hypothalamic or a pituitary reason as a cause of the menstrual irregularities. The ICF, which predicted a 66% probability, would prompt the individual to seek advice from a medical professional via the Positive Result Screen, and also suggest PCOS as a possible cause for signs and symptoms reported.

For virtual test subject 12, the physician assigned a 10% probability of PCOS. The ICF also suggested a 10% probability of PCOS. The physician determined that the hormones taken by the individual indicated a special case, and that recommendations could not be made based on the answers provided. The ICF’s Inconclusive Screen was presented to this virtual test subject.

For virtual test subject 15, both the physician and the ICF predicted a 0% to 1% chance of PCOS and the subject received the Neutral Result Screen and Doctor’s Report reporting PCOS is an unlikely cause of the menstrual irregularities.

## Discussion

The team combined the expertise of data scientists, software engineers, and a medical expert to construct a mobile health tool that draws attention to possible indicators of PCOS in an app using population. To the best of our knowledge, this is the first app developed by an interdisciplinary team to calculate the probability that an individual may have a risk of PCOS. Furthermore, the ICF allows users to self-report menstrual information to facilitate a discussion with their doctors. Because a summary generated based on the answers to the interactive survey is specific to each individual, more control is in the hands of the user. Additionally, by providing the user with an outline of what the next steps are in terms of testing and visiting a doctor’s office, upon roll-out to Clue users, this novel tool could shorten the time for a PCOS diagnosis.

As seen in virtual test subject 1, the similarly high probabilities assigned by the physician and ICF demonstrate that in textbook cases of PCOS, the ICF accurately prompts individuals to seek out a medical professional. This will be important for those indicating several risk factors for PCOS. Virtual test subject 8 highlights the over predictive nature of the ICF. Despite the ICF probability being slightly higher than the physician’s prediction, the tool still proves useful as it advises an individual to seek a healthcare provider for further testing. Additionally, the lack of data input for virtual test subject 8, specifically for menstrual variation and confounding diagnoses, can be improved once data is collected from actual Clue® users in order to make more accurate predictions. Virtual test subject 12, who is using hormonal birth control, demonstrates how the ICF calculates a probability for individuals inputting the minimum amount of information and confounders while also indicating menstrual irregularities. The ICF suggests similar individuals to seek a medical professional for any issues regarding their menstrual irregularities. Virtual test subject 15 did not report menstrual irregularities nor did they indicate areas of concern such as those associated with hyperandrogenism: acne, alopecia, or hirsutism. This illustrates that the model can accurately eliminate individuals who are unlikely cases of the disorder.

The main limitations of this study are that it only used a small number of virtual test subjects, was not tested in a human population, and that the probability generated by the tool is not a diagnosis. Additionally, the model cannot make predictions for individuals who report use of hormone-based medications or other syndromes similar to PCOS, such as Cushing’s syndrome or hypothyroidism. The ICF also errs on the side of sending more individuals to the doctor’s office as an attempt to capture the majority of individuals at risk for PCOS. It remains to be seen whether the predictive model will be adaptable to actual human users. A future validation study of the ICF will be needed to assess the utility of the tool by following up with users at variable time intervals in order to determine if a medical professional diagnosed PCOS or another condition.

Despite the limitations of the study, the ICF is one example of how mobile technology may help users manage their own health and promote subject self-advocacy. By providing more information about PCOS and giving users feedback based on information they have collected, the research team predicts that this feature will provide important information to those at high risk of the disease and potentially shorten the time for diagnosis. In the future, the team anticipates that this may be a valuable tool for doctors and subjects alike. The creation of the Irregular Cycles Feature may reduce the time for PCOS diagnosis and facilitate conversations between users and doctors through the Results Screens and Doctor’s Report.

## Acknowledgements

The research team would like to acknowledge Pascaline Karanja, MS for proofreading the article and Virginia Vitzthum for supporting the collaboration at Clue.

