## Supplementary Figures for "Identifying Women at Risk for Polycystic Ovary Syndrome Using a Mobile Health Application"

**Supplementary Figure 1. Image of Positive Result Screen**

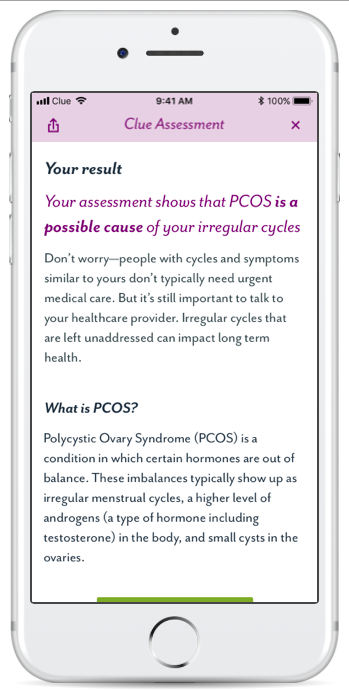

**Supplementary Figure 2. Doctor’s Report for PCOS as a likely cause of menstrual irregularity**

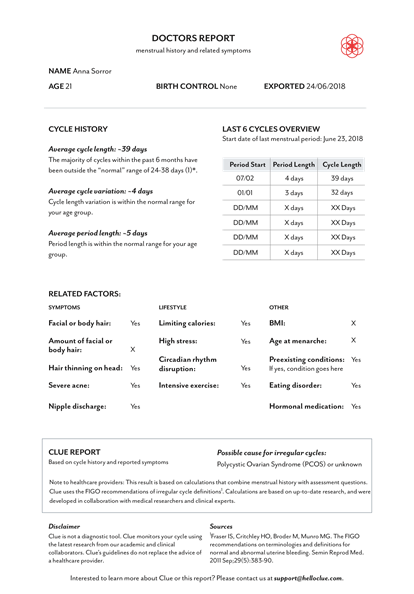

**Supplementary Figure 3. Image of Clue Graphic User Interface**

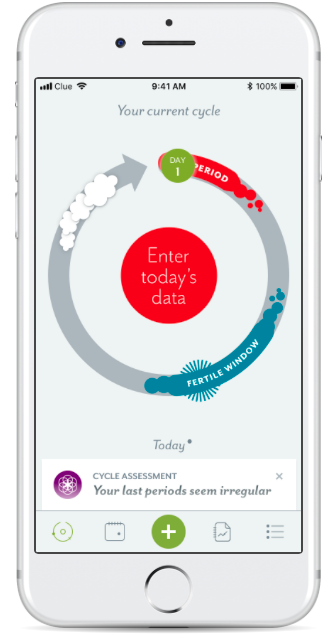

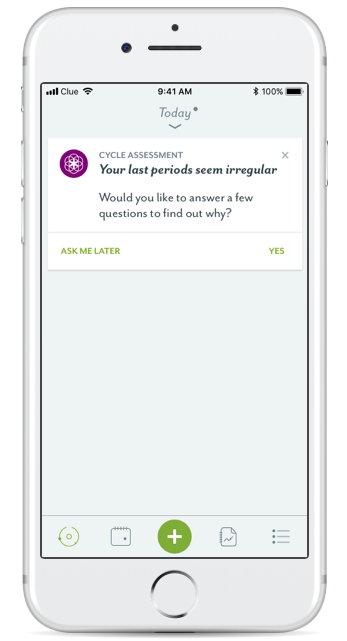
